# The N-terminus of GPR37L1 is proteolytically processed by matrix metalloproteases

**DOI:** 10.1101/2020.06.27.174847

**Authors:** James L. J. Coleman, Tony Ngo, Rhyll E. Smythe, Andrew J. Cleave, Nicole M. Jones, Robert M. Graham, Nicola J. Smith

## Abstract

GPR37L1 is an orphan G protein-coupled receptor expressed exclusively in the brain and linked to seizures, neuroprotection and cardiovascular disease. Based upon the observation that fragments of the GPR37L1 N-terminus are found in human cerebrospinal fluid, we hypothesized that GPR37L1 was subject to post-translational modification. Heterologous expression of GPR37L1-eYFP in either HEK293 or U87 glioblastoma cells yielded two cell surface species of approximately equivalent abundance, the larger of which is N-glycosylated at Asn^105^. The smaller species is produced by matrix metalloprotease/ADAM-mediated proteolysis (shown by the use of pharmacological inhibitors) and has a molecular weight identical to that of a mutant lacking the entire N-terminus, Δ122 GPR37L1. Serial truncation of the N-terminus prevented GPR37L1 expression except when the entire N-terminus was removed, narrowing the predicted site of N-terminal proteolysis to residues 105-122. Using yeast expressing different G protein chimeras, we found that wild type GPR37L1, but not Δ122 GPR37L1, coupled constitutively to Gpa1/Gαs and Gpa1/Gα16 chimeras, in contrast to previous studies. We tested the peptides identified in cerebrospinal fluid as well as their putative newly-generated N-terminal ‘tethered’ counterparts in both wild type and Δ122 GPR37L1 Gpa1/Gαs strains but saw no effect, suggesting that GPR37L1 does not signal in a manner akin to the protease-activated receptor family. We also saw no evidence of receptor activation or regulation by the reported GPR37L1 ligand, prosaptide/TX14A. Finally, the proteolytically processed species predominated both *in vivo* and *ex vivo* in organotypic cerebellar slice preparations, suggesting that GPR37L1 is rapidly processed to a signaling-inactive form. Our data indicate that the function of GPR37L1 *in vivo* is tightly regulated by metalloprotease-dependent N-terminal cleavage.

## Introduction

G protein-coupled receptors (GPCRs) are the largest family of integral membrane proteins encoded by the human genome, with over 800 unique members (1). They are a major class of pharmacological targets – a function of the high degree of receptor-ligand specificity, the extracellular localization of their ligand-binding sites, and the discrete distribution of individual GPCRs in the body (2). Indeed, an estimated 34% of existing, marketed pharmaceuticals act at GPCRs (3). These receptors are typically activated by binding of one or more of a diverse range of extracellular stimuli, including photons, peptide hormones, lipids, amines, and ions, leading to activation of heterotrimeric G proteins and subsequent signal amplification (4). Depending upon the receptor conformation stabilized by the ligand, multiple distinct pathways can be activated or repressed. To maintain their exquisite spatial and temporal specificity, GPCRs are tightly regulated by post-translational modifications that affect their trafficking to the cell surface, internalization and down-regulation (5).

The GPR37-like 1 receptor, or GPR37L1, is an orphan GPCR (no identified endogenous partner) expressed diffusely in glial cells of the central nervous system, with particularly high expression in Bergmann glia and astrocytes in the cerebellum (6-9). Mice lacking GPR37L1 have been reported to have precocious cerebellar development and motor learning (10), although this has been disputed (9), as well as increased susceptibility to seizures (11) and chemical ischemia (9). In the periphery, we and others have reported that mice lacking GPR37L1 have cardiovascular phenotypes of varying severity (8,12), however the lack of detectable GPR37L1 protein in murine heart or kidney suggests this effect is also CNS-mediated (8). Most recently, GPR37L1 has also been linked to oncogenesis as it was found to be over-transcribed in human glioblastomas (13), while reduced expression in medulloblastomas may be responsible for tumour progression (14). These initial reports suggest that GPR37L1 may play a role in various physiological and pathophysiological processes, yet little is known about the receptor itself.

Although GPR37L1 shares 32% identity and 56% similarity with the endothelin-B receptor (6,7), it does not bind endothelin or related peptides (7). Its closest relative, GPR37 (parkin-associated endothelin-like receptor), is also an orphan. Currently, the molecular pharmacology of GPR37L1 is poorly understood (15), although the neurotrophic factor prosaposin [and its synthetic peptide derivative prosaptide (TX14A)] have been proposed as endogenous GPR37L1 and GPR37 ligands (16). Intriguingly, analyses of normal human cerebrospinal fluid (CSF) identified peptides (17-19) that are identical to three distinct regions of the GPR37L1 N-terminus, suggesting that perhaps GPR37L1 is processed and activated by a mechanism akin to that of the protease-activated receptors (PARs) (20). In this study^*(See Footnote)*^, we tested the hypothesis that the N-terminus of GPR37L1 is post-translationally modified and that this may influence receptor function. We report that GPR37L1 is subject to metalloprotease-dependent proteolytic processing of its N-terminus, both when expressed in cultured cells and in rodent cerebellar tissue. Using a yeast G protein chimera assay (21,22), we found that GPR37L1, but not a mutant lacking the entire N-terminus, can signal via Gαs but that, unlike PARs, synthetic peptides corresponding to the N-terminus of GPR37L1 did not activate the receptor. These findings indicate that GPR37L1 signaling is negatively regulated by metalloprotease-mediated N-terminal processing.

## Results

### GPR37L1 is N-terminally processed

We stably transfected human GPR37L1 constructs in which the C-terminus is fused to enhanced yellow fluorescent protein (GPR37L1-eYFP) into FlpIN T-REx HEK293 cells. We have previously used this doxycycline-inducible system to evaluate orphan or low affinity GPCRs because it provides, in the absence of induction, negative expression controls that are essential when working with poorly characterized receptors (23-28). We generated serial N-terminal truncations to assess the effect of removing the putative signal peptide (Δ25 GPR37L1-eYFP), the signal peptide plus the region encompassing all three reported N-terminal fragments found in CSF (Δ80 GPR37L1-eYFP), or the entire N-terminus (Δ122-GPR37L1-eYFP), on receptor production and delivery to the cell surface (**Figure 1a**).

**Figure 1.**
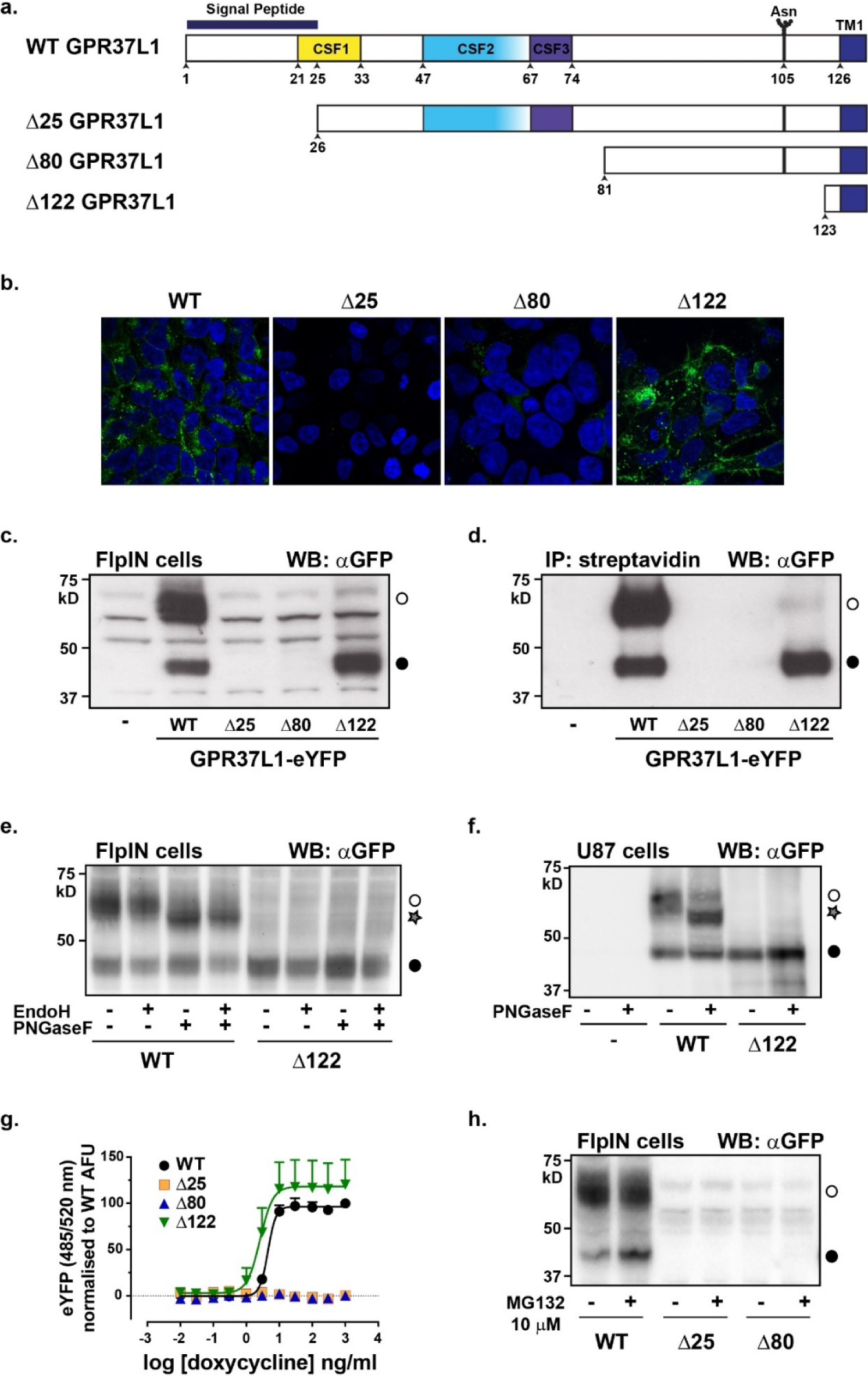
GPR37L1 exists as both a full-length and N-terminally truncated species. **(a)** Schematic of the GPR37L1 N-terminus and location of serial truncations. CSF1-3: cerebrospinal fluid-derived peptides identified by mass spectrometry (17-19). Asn^105^ is the predicted glycosylation site. TM1: start of transmembrane domain 1. **(b)** Confocal microscopy (x63 magnification) of each FlpIN T-REx HEK293 GPR37L1-eYFP stable cell line induced with 1 μg/ml doxycycline for 24 h. Merged image of eYFP fluorescence (receptor) and Hoechst 33342 (nucleus). **(c)** Immunoblot of lysates from each stable cell line with an antibody targeted to GFP. Cells were induced with 500 ng/ml doxycycline for 24 h before harvest. **(d)** Immunoprecipitation of biotinylated lysates shown in (c). Open circle, full length receptor. Closed circle, truncated receptor. **(e)** FlpIN stable cells induced for 24 h with 500 ng/ml doxycycline were lysed and treated for 2 h with either Endoglycosidase H (EndoH, 100 mU) or peptide-*N*-glycosidase F (PNGaseF, 2U), or both, before immunoblotting with the GFP antibody. Star symbol represents deglycosylated receptor. **(f)** U87 human glioblastoma cells transiently expressing GPR37L1 or Δ122 GPR37L1-eYFP were treated as per (e). **(g)** Whole cell eYFP fluorescence (excitation 485 nm, emission 520 nm) in response to increasing concentrations of doxycycline; AFU: arbitrary fluorescence units. **(h)** Immunoblot from cells treated for 6 hours with proteasome inhibitor MG132 (10 μM) before lysis. Data are representative of three independent experiments except for **(g)**, which shows the mean values +/- S.E.M. for n = 5.

Induction of wild-type GPR37L1-eYFP with doxycycline resulted in apparent cell surface expression and robust accumulation of the receptor (**Figures 1b & 1c**). Of note, GPR37L1-eYFP resolved as two molecular weight species by SDS-PAGE; a larger band of M_r_ ∼65-70 kD (**Figure 1c**, open circle) and a smaller band of ∼40-45 kD (closed circle). The smaller molecular weight species was equivalent in size to Δ122 GPR37L1-eYFP, which we detected as a single species. Cell surface biotinylation labeled both GPR37L1-eYFP species and the relative ratio of larger molecular weight to smaller molecular weight species appeared similar between whole-cell lysates and biotinylated samples (**Figure 1c vs 1d**). We also detected Δ122 GPR37L1-eYFP at the cell surface, thus we reasoned that the smaller species of GPR37L1 might represent an N-terminally processed version of the receptor. Deglycosylation of the single predicted glycosylation site, Asn^105^ (6), failed to reduce the larger band to the same size as the smaller band (**Figure 1e**), indicating that the smaller species is unlikely to be a precursor of full-length GPR37L1, which is consistent with the biotinylation results. Neither the smaller species, observed in cells expressing GPR37L1-eYFP, nor the species in cells expressing Δ122 GPR37L1-eYFP, showed a decrease in molecular weight in response to deglycosylation, consistent with cleavage of the N-terminus occurring after Asn^105^. Furthermore, we could exclude C-terminal truncation because both the full-length and truncated versions of GPR37L1-eYFP were visible by in-gel fluorescence (data not shown and **Figure 2a**), indicating that the C-terminal eYFP fusion was still present and correctly folded. Finally, N-terminal cleavage of GPR37L1-eYFP was not unique to HEK293 cells, as the same GPR37L1-eYFP processing and response to deglycosylation was also observed in the more physiologically-relevant human glioblastoma U87 MG cell line transiently transfected with either wild-type or Δ122 GPR37L1-eYFP (**Figure 1f**).

**Figure 2.**
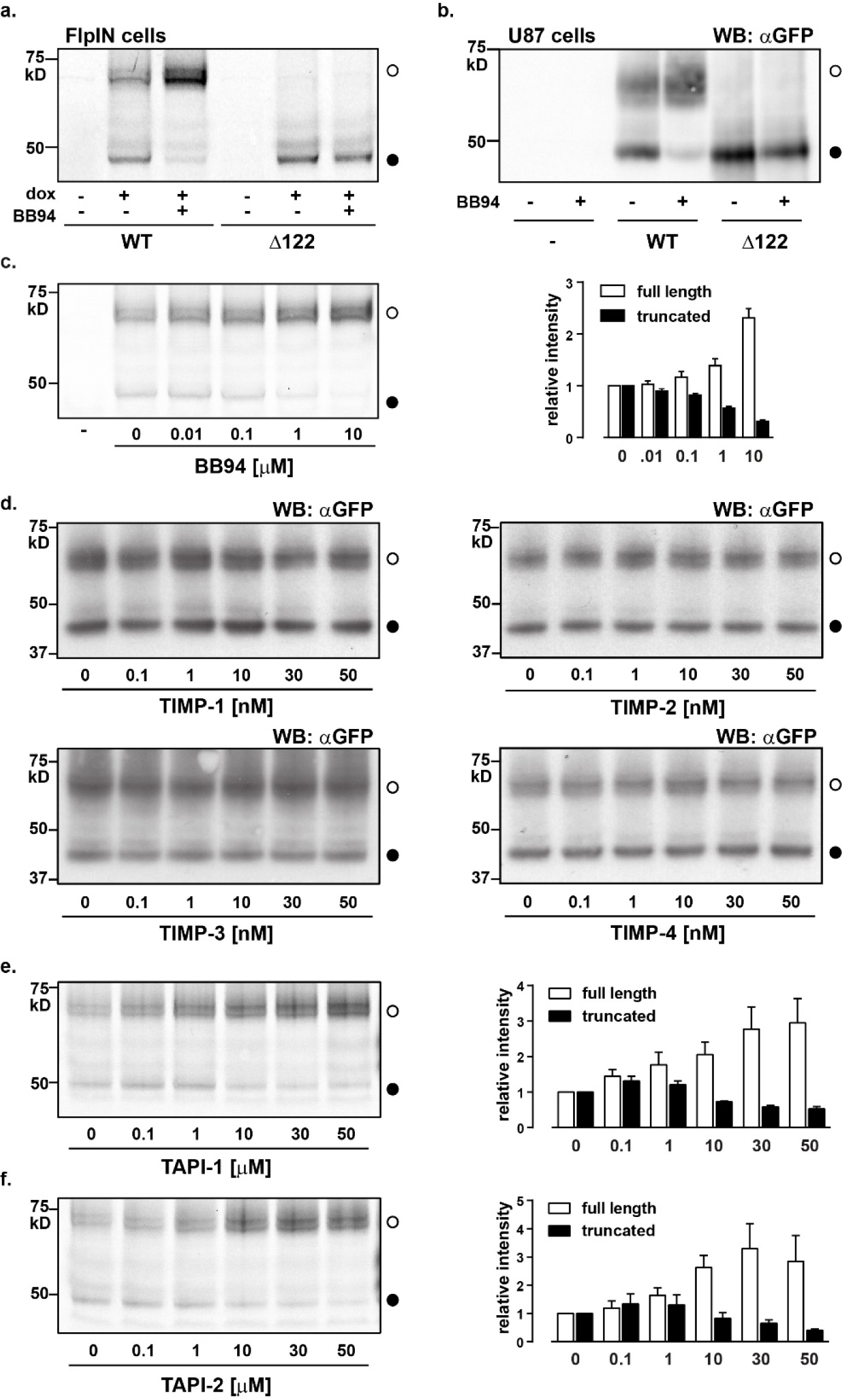
GPR37L1 is processed at the N-terminus by an ADAM. FlpIN HEK293 stable cell lines were used for all experiments except for (b) and were induced with 500 ng/ml doxycycline. **(a)** Wild-type (WT) or Δ122 GPR37L1-eYFP expression was induced in the presence of BB94 (20 µM) for 24 h and visualized by in-gel fluorescence. **(b)** U87 glioblastoma cells were transiently transfected with WT or Δ122 GPR37L1-eYFP and treated for 24 h with 20 µM BB94 before immunoblotting for fluorescent protein expression. **(c)** GPR37L1-eYFP cells were induced in the presence of increasing concentrations of BB94 for 24 h (quantified in the right panel). ‘-’ = uninduced. Protein was visualized by in-gel fluorescence. **(d)** GPR37L1-eYFP cells were induced in the presence of increasing concentrations of human recombinant TIMP-1, TIMP-2, TIMP-3, or TIMP-4 for 24 h and then immunoblotted with a GFP antibody. Biological activity of recombinant TIMPs was validated using activated MMP2 and a fluorogenic substrate (**Supplementary Figure 1**). AFU, arbitrary fluorescence units. **(e & f)** GPR37L1-eYFP cells were induced in the presence of the indicated concentrations of TAPI-1 or TAPI-2 for 24 h and the amount of full-length and truncated species were quantified compared to untreated cells. Left panel: In-gel fluorescence of cell lysates; Right panel: quantification of full-length (∼65-70 kD, open circle) and truncated (∼40-45 kD, closed circle) GPR37L1 fluorescence. Each image is representative of at least four individual experiments.

Unlike the Δ122 GPR37L1-eYFP construct, which presumably reaches the cell surface as a result of a cryptic signal sequence within its first transmembrane domain (29), we could not detect either the Δ25 or Δ80 truncated forms, even when we exposed the cells to the highest concentrations of doxycycline (1 µg/ml), as confirmed by confocal microscopy (**Figure 1b**), immunoblotting (**Figure 1c and 1d**) and whole-cell eYFP fluorescence (**Figure 1g**). To determine if this was due to rapid degradation of improperly folded receptor, we exposed cells for 6 hours to the proteasomal degradation inhibitor, MG132 (Z-Leu-Leu-Leu-H). However, the proteins were undetectable under these conditions (**Figure 1h**), indicating that these truncated versions could not be properly folded or synthesized.

### The N-terminus of GPR37L1 is truncated by a metalloprotease

Because some GPCRs undergo metalloprotease-catalyzed N-terminal cleavage (30-32), we hypothesized that MMPs (matrix metalloproteases) or ADAMs (a disintegrin and metalloproteases) cleave GPR37L1. Induction of GPR37L1 expression in the presence of a single concentration of the nonspecific MMP and ADAM inhibitor, BB94 (batimastat), almost completely prevented the generation of the smaller receptor species in cells expressing GPR37L1-eYFP (**Figure 2a**). The same effect was observed in U87 glioblastoma cells (**Figure 2b**). Furthermore, we observed a concomitant concentration-dependent increase in full-length GPR37L1-eYFP and reduction in the smaller species (**Figure 2c**), consistent with a product-precursor relationship between the two proteins. The presence of BB94 did not alter the abundance of Δ122 GPR37L1-eYFP, which is expected because this form already lacks the N-terminus.

Tissue inhibitors of metalloproteases (TIMPs) are endogenous inhibitors of both MMPs and ADAMs and have overlapping but not identical inhibitory profiles for their substrates. TIMP-1 is an inhibitor of ADAM10 and shows weak inhibition of MMP-14, MMP-16, MMP-19, and MMP-24. TIMP-2 inhibits all MMPs and also ADAM12. TIMP-3 similarly inhibits all MMPs and also ADAM10, ADAM12, ADAM17, ADAM28, and ADAM33. TIMP-4 also displays a wide MMP inhibitory profile with further inhibition at ADAM17, ADAM28, and ADAM33 (33). GPR37L1-eYFP induced in the presence of 4 different TIMPs still produced both the full-length and short forms (**Figure 2d**). The lack of an effect was not due to inactivity of the recombinant TIMPs, because the same batch of each recombinant TIMP inhibited recombinant MMP2 (**Supplementary Figure 1**). These data indicated that all MMPs as well as ADAM10, 12, 17, 28, and 33 were unlikely mediators of GPR37L1 N-terminal cleavage.

ADAMs and MMPs with a Zn^2+^-containing catalytic domain can be inhibited by nonspecific inhibitors that interact with the catalytic domain. These inhibitors include BB94 and tumor necrosis factor-α protease inhibitors-1 (TAPI-1) and −2 (TAPI-2) (34). Like the results with BB94 (**Figure 2c**), the presence of TAPI-1 or TAPI-2 prevented GPR37L1-eYFP cleavage in a concentration-dependent manner (**Figure 2e & 2f**). From these data, we conclude that one or more ADAM family members that are not inhibited by TIMPs but have a Zn^2+^ catalytic domain, that is ADAMs 8, 9, 15, 19, 20, 21, and 30 (35), may be responsible for GPR37L1 N-terminal proteolysis.

### GPR37L1 N-terminal proteolysis is preserved following alanine mutagenesis

We reasoned that GPR37L1 must be cleaved between the predicted glycosylation site (Asn^105^) and the beginning of transmembrane 1 (after residue 122), so we performed alanine scanning mutagenesis in triplets to identify the site of cleavage (**Figure 3a**). Except for ^109^Ala^111^ GPR37L1-eYFP, which could not be detected by immunoblot, all mutants expressed two species of GPR37L1-eYFP (**Figure 3b**). Deglycosylation reduced the apparent molecular weight of all mutants except for ^106^Ala^108^ GPR37L1-eYFP, which has lost its NxS/T N-linked glycosylation motif (**Figure 3c**). N-terminal proteolysis was not prevented by any mutation and each receptor remained sensitive to metalloprotease inhibition (**Figure 3d**). These findings are consistent with the notion that metalloproteases lack clearly defined consensus sequences (36) or that the non-expressing ^109^Ala^111^ species may indeed have had perturbed proteolysis.

**Figure 3.**
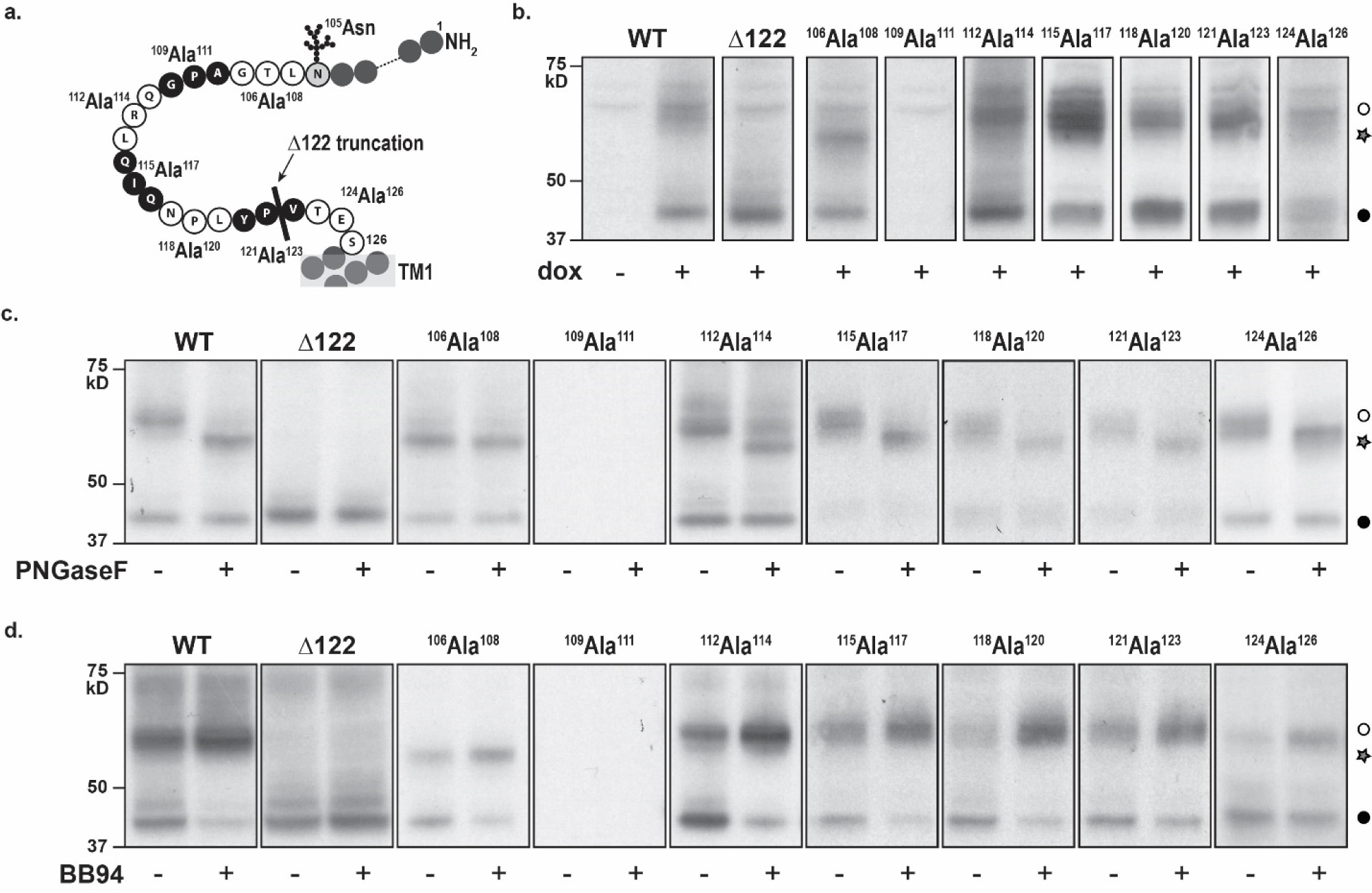
Systematic triplet alanine scanning mutation reveals residues necessary for GPR37L1-eYFP glycosylation and expression. Stable cell lines of inducible GPR37L1-eYFP triplet alanine mutants, indicated in the schematic **(a)**, were assessed for effect on **(b)** expression, **(c)** glycosylation status, and **(d)** sensitivity to metalloprotease inhibition. PNGase F = 2 U, BB94 = 10 μM. TM1: transmembrane domain 1. Open circle: full-length receptor. Closed circle, truncated receptor. Star, deglycosylated GPR37L1. n = 4.

### GPR37L1 constitutively couples to chimeric yeast Gpa1/Gαs

To understand the biological relevance of N-terminal processing of GPR37L1, we sought to identify the signal transduction pathways activated by this receptor. We employed a G protein coupling screen using *Saccharomyces cerevisiae* with β-galactosidase as the output, which enabled assessment of GPR37L1 activity without prior knowledge of its ligand or signal transduction pathway. In this system, the candidate receptor is transformed into an engineered yeast strain with a modified pheromone-response pathway, where the yeast Gα subunit, Gpa1, has had its final five amino acids replaced by the corresponding sequence for each of the human Gα species. As such, receptor activity transmitted through the modified heterotrimeric G protein will lead to dissociation of the native Gβγ dimer, which in turn triggers a mitogen-activated protein kinase cascade and ultimately results in transcription of a β-galactosidase reporter gene driven by a FUS1 promoter (21,22). We have previously used this system to identify the G protein coupling profile of another orphan receptor, GPR35 (23).

In this system, GPR37L1-eYFP coupled to the chimeric G protein, Gpa1/Gαs as well as Gpa1/Gα16, but not the other Gαq/11 family chimeras (**Figure 4a**). Moreover, we did not observe any constitutive coupling to members of the Gαi/o family, nor did the receptor couple to either Gα12 or Gα13 (**Figure 4a**). In contrast to the full-length receptor, Δ122 GPR37L1-eYFP did not signal through any of the tested G protein chimeras (**Figure 4a**). Given that Gα16 (now known as Gα15 in humans) couples promiscuously to GPCRs that signal through the Gαs, Gαi/o, or Gαq/11, but not Gα12/13, G protein families (37,38), these findings suggest that GPR37L1 may be a Gαs- and, thus, adenylyl cyclase-coupled receptor.

**Figure 4.**
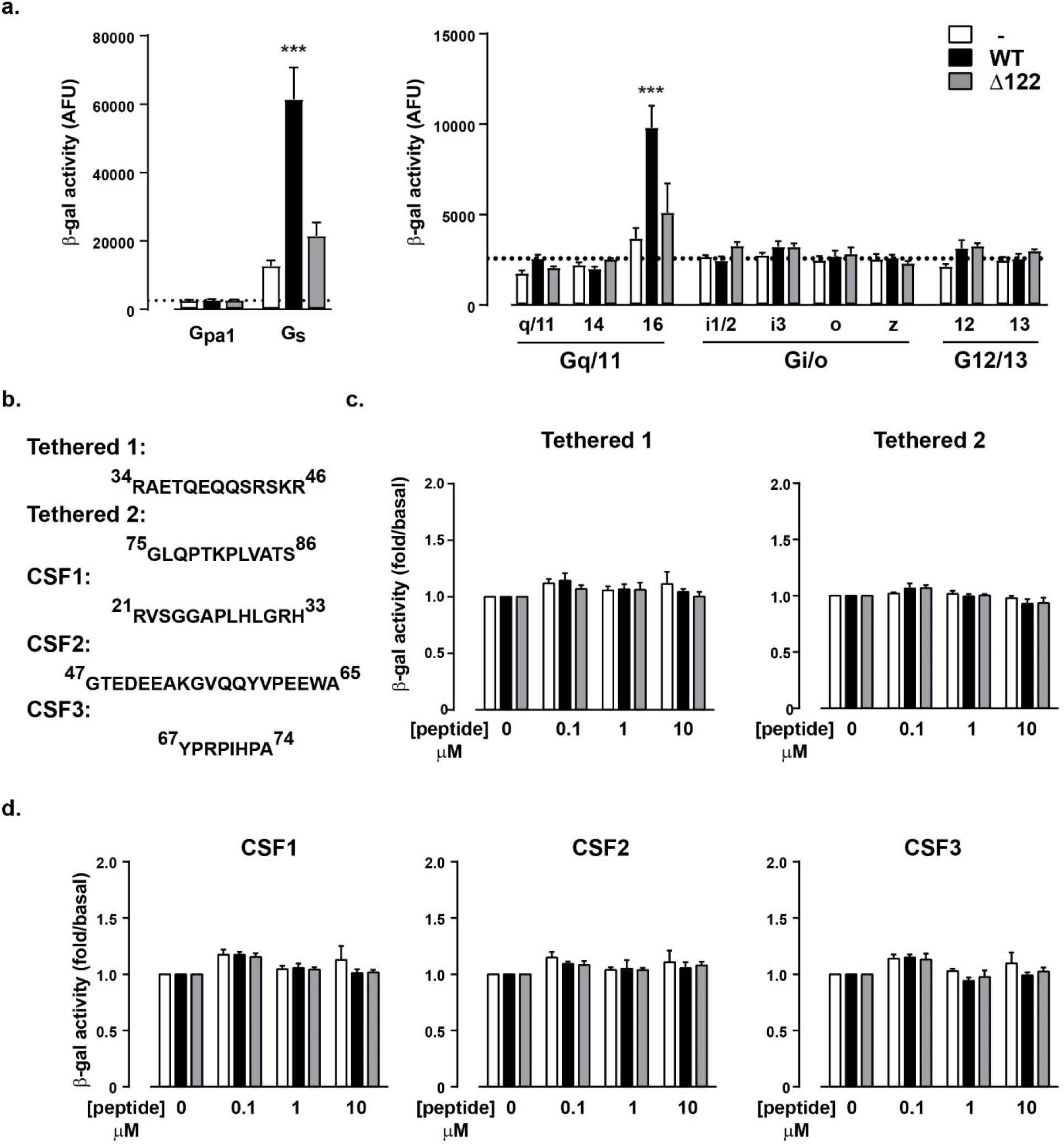
GPR37L1 N-terminus is necessary for Gαs constitutive activity in yeast but is not auto-activating. **(a)** WT GPR37L1-eYFP, Δ122 GPR37L1-eYFP, or empty p426GPD vector was transformed into yeast strains bearing individual G protein chimeras from each of the four G protein families (22), and β-galactosidase (β-gal) activity was measured in arbitrary fluorescence units (AFU) after 24 hours. Dotted line represents vector-transformed Gpa1 β-galactosidase activity for comparison. n = 9 individual transformants. **(b)** Synthetic peptides with corresponding GPR37L1 sequence numbers indicated were tested (3 individual transformants) for Gαs-mediated β-galactosidase activity. **(c)** Putative tethered peptides, tethered 1 and tethered 2. **(d)** CSF peptides identified from mass spectrometry (17-19). \*\*\**p*<0.001 when compared to vector alone, according to a one-way ANOVA with Bonferroni’s post-hoc analysis.

Because the detection of N-terminal peptides derived from GPR37L1 in CSF indicated that, like PARs, GPR37L1 is proteolytically processed, we examined if short peptides corresponding to regions identified as proteolytic products of GPR37L1 in human CSF (17-19) could activate the receptor. We call the three peptides CSF1 (residues 21-33), CSF2 (residues 47-65), and CSF3 (residues 67-74) (**Figure 1a & Figure 4b**). Cleavage of the N-terminus to release CSF1 (residues 21-33) would leave the N-terminal domain from residue 34 to the first transmembrane domain at residue 126. Within this remaining N-terminal section would be sequences that produce CSF2 (residues 47-65) and CSF3 (residues 67-74). Cleavage to release CSF2 would eliminate the N-terminal region to residue 68, just at the position where CSF3 commences. Finally, proteolytic cleavage of CSF3 would eliminate all three peptides from the N-terminal region. We also tested if addition of the peptides corresponding to residues 34-46 (“tethered-1”, the N-terminal end up to the CSF2 sequence that would be produced by cleavage and release of CSF1) or residues 75-86 (“tethered-2”, an N-terminal peptide sequence of arbitrary length that would be revealed upon cleavage and release of CSF3), altered receptor activity. However, we found no effect of the CSF peptides or tethered peptides in the yeast chimeric G protein assay when tested with wild-type GPR37L1-eYFP or Δ122 GPR37L1-eYFP (**Figure 4c & d**).

### GPR37L1 is N-terminally processed in vivo

Thus far, our evidence for proteolytic processing of the N-terminus has come from heterologously expressed GPR37L1-eYFP in FlpIN HEK293 or human glioblastoma cells. To confirm that this processing occurs *in vivo* and is physiologically-relevant, we examined GPR37L1 abundance in the cerebellum from C57BL/6J wild-type and GPR37L1-null mice (12). We chose cerebellum because GPR37L1 mRNA and protein is most abundant in this tissue and global GPR37L1-knockout mice are reported to have a potential cerebellar phenotype (6,7,9,10). Furthermore, there is little difference in sequence between orthologs, with mouse and rat sharing 91% identity and 100% similarity with human GPR37L1. In the cerebellum of male mice, we found that GPR37L1 existed almost entirely as the smaller molecular weight species in this tissue (**Figure 5a**). No GPR37L1 was identified in the GPR37L1-null tissue, confirming that the antibody used to detect GPR37L1 was specific for the receptor. Based upon our yeast experiments in Figure 4a, the ratio of smaller to larger MW GPR37L1 would suggest that receptor signaling may be silenced in this tissue.

**Figure 5.**
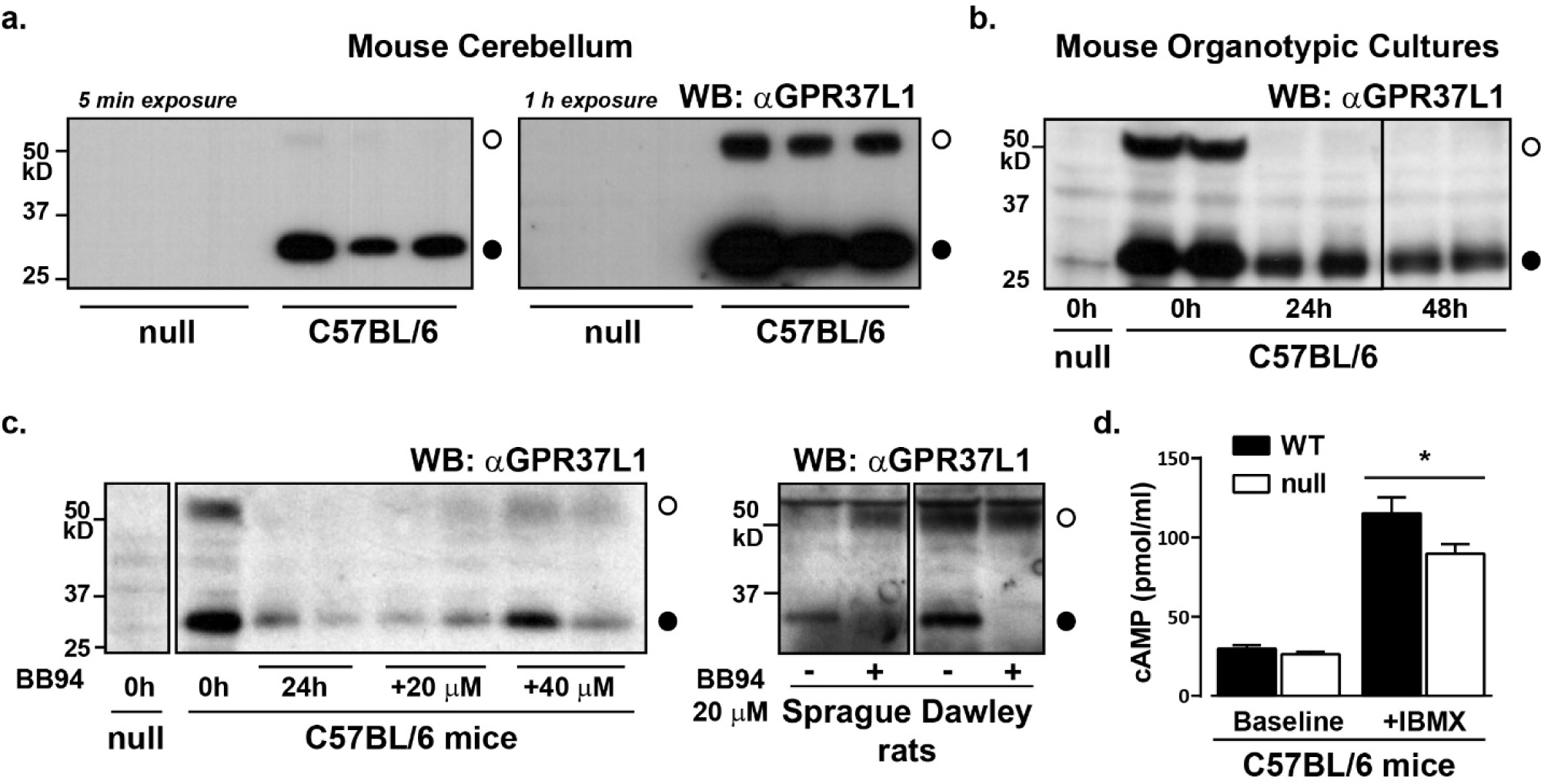
GPR37L1 is N-terminally proteolytically processed in vivo. **(a)** Plasma membrane was isolated from the cerebella from adult GPR37L1-null or C57BL/6J male mice and samples probed for endogenous GPR37L1. n = 3. The remaining experiments were performed using mouse or rat organotypic cultures, as indicated. **(b)** Mouse cerebella were isolated from either wild-type C57BL/6J or GPR37L1-null 15-day-old mixed-sex mice and slice preparations were cultured for varying times before harvest and membrane preparation. n = 2 as shown. **(c)** Separate mouse or rat cerebellar slice preparations were immediately exposed to 20 µM or 40 µM BB94 for 24 h (mouse), or to 20 µM BB94 after 10 days in culture for 48 h (rat, 8-9 days old, mixed-sex), before measurement of GPR37L1 expression. Shown is n = 2 of 4 (mouse) or 6 (rat) experiments. **(d)** cAMP accumulation in cerebellar slices from C57BL/6J (WT) or GPR37L1-null mice were cultured for 1 h in the presence or absence of 1 mM IBMX. n = 6, where each sample contains tissue from 2-4 separate mice. *p<0.05.

To assess cAMP generation and ADAM- or MMP-dependent cleavage in an endogenous setting, we established cerebellar organotypic slice preparations. Immediately after isolation, the cultured tissue had both full-length and the shorter form of GPR37L1 (**Figure 5b**). We found that the abundance of both species of GPR37L1 was markedly reduced within 24 h of culture (**Figure 5b**). Despite this, we were able to detect the appearance of full length GPR37L1 if cerebellar preparations were immediately placed into BB94 MMP inhibitor-containing medium for 24 h (mouse) or 48 h (rat) (**Figure 5c**). Together, these results showed that a metalloprotease inhibitor reduced the processing of GPR37L1 in both rat and mouse cerebellum.

Finally, because we had observed Gαs coupling in the chimeric yeast assay, we examined the functional consequence of GPR37L1 deletion by measuring cerebellar cAMP concentrations in cultured slices by ELISA, either in the absence or presence of phosphodiesterase inhibition with IBMX. Although baseline cAMP accumulation was the same between C57BL/6J and GPR37L1-null mice, incubation of the slices with IBMX for one hour revealed that deletion of GPR37L1 resulted in significantly less cAMP production in the cerebellum (**Figure 5d**). Thus, despite the observation that GPR37L1 exists predominantly as a processed, and potentially inactive species, mice lacking GPR37L1 have less cAMP production in the cerebellum. Thus, either the remaining full-length GPR37L1 is sufficient to account for the observed difference in cAMP, or the proteolytically processed form of GPR37L1 retains the capacity to signal *in vivo*.

### GPR37L1 does not bind prosaptide (TX14A)

A study identified the neurotrophic synthetic peptide prosaptide (TX14A), which is derived from the saposin C cleavage product of the endogenous prosaposin protein, as a dual GPR37 and GPR37L1 agonist that induces receptor internalization and Gαi–mediated signal transduction (16). We tested the effect of TX14A on GPR37L1-mediated G protein coupling in the yeast chimeric G protein assay for Gαs, which we found coupled to GPR37L1 (**Figure 4a**), and for Gαi1 and Gαi3, the G proteins reportedly stimulated by TX14A (16). GPR37L1-eYFP coupling to the Gαs chimeric protein was the same in the presence or absence of TX14A, and TX14A did not stimulate coupling to either of the Gαi chimeric subunits (**Figure 6a**). To rule out the possibility that the purchased TX14A was spoiled or inactivated, we tested multiple batches of TX14A with the same outcome. These data indicate GPR37L1 does not signal in response to TX14A in the yeast G protein coupling assay.

**Figure 6.**
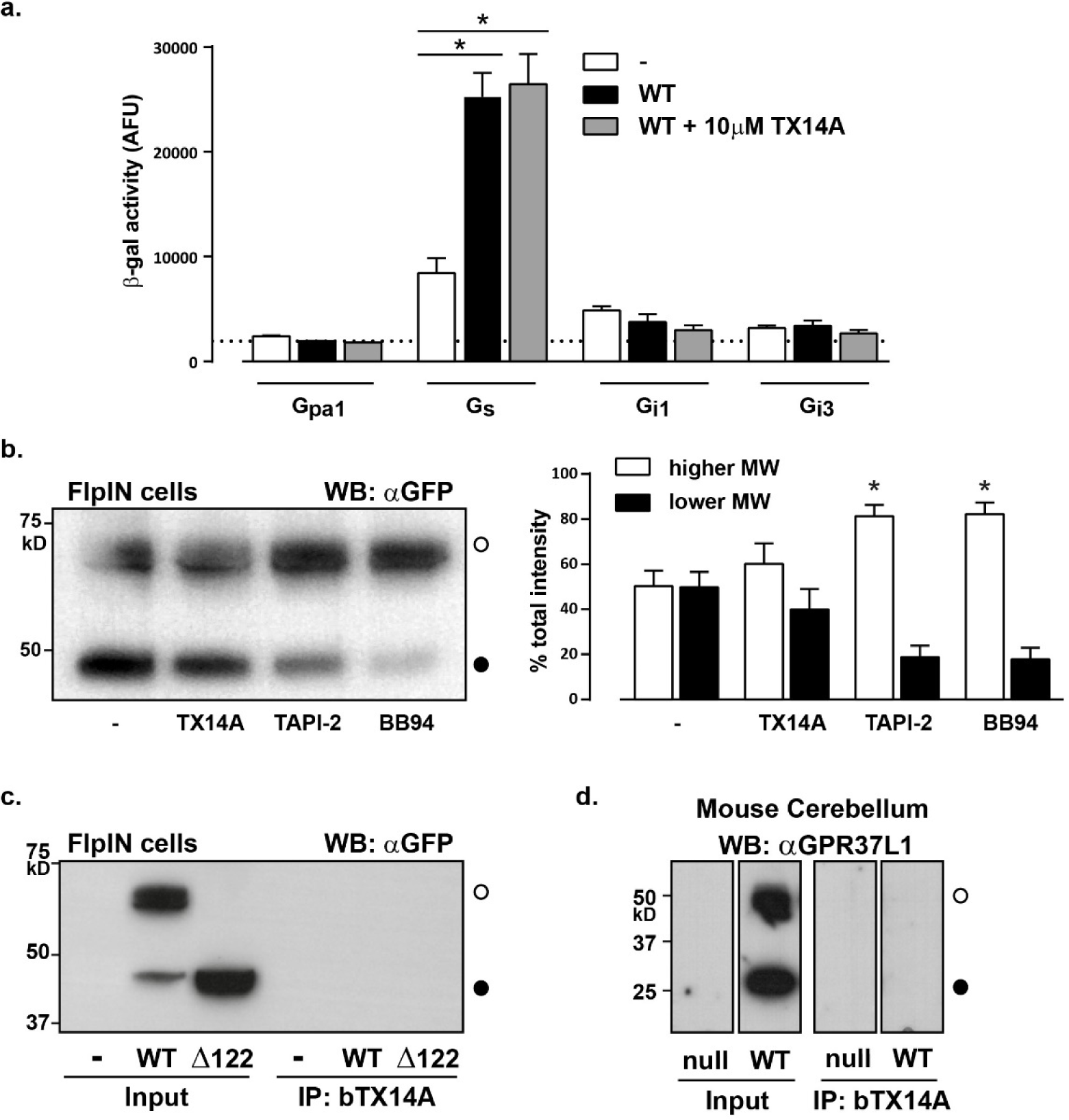
TX14A does not stimulate GPR37L1 signaling or alter protein abundance. **(a)** β-galactosidase activity in Gpa1 or Gαs, Gαi1 or Gαi3 yeast chimera strains transformed to express either p426GPD vector or WT GPR37L1-eYFP. Transformants were stimulated with 10 µM TX14A for 24 h. n = 12 individual transformants per strain. **(b)** FlpIN cells were induced with doxycycline (500 ng/ml) for 24 h to express WT GPR37L1-eYFP and then exposed to 10 µM TX14A, 10 µM TAPI-2, or 20 µM BB94 for 24 h. A representative blot is shown on the left and on the right is densitometry of the relative GPR37L1 species as a percentage of total GPR37L1 (both species). n = 4. **(c)** Analysis of TX14A binding to membranes from FlpIN cells expressing WT or Δ122 GPR37L1-eYFP. Membrane preparations were incubated with biotinylated TX14A (bTX14A) and then precipitated (IP) with streptavidin beads. Input and immunoprecipitated samples were probed with an antibody for GFP. Representative blot, n = 3. **(d)** Analysis of bTX14A binding to cerebellar membranes from wild-type or GPR37L1-null mice, as per (c). Samples were probed with an antibody raised against GPR37L1, n = 3. *p<0.05 according to a one-way ANOVA with Bonferroni’s post-hoc analysis.

We next determined whether chronic treatment might reveal an effect of TX14A at the level of protein abundance. Both TAPI-2 and BB94 significantly increased the ratio of full-length to truncated GPR37L1-eYFP without altering the overall amount of the receptor (as per Figure 2), whereas TX14A treatment did not significantly change this ratio or apparent abundance (**Figure 6b**). Finally, in the absence of evidence for a signaling or regulatory role for TX14A, we aimed to confirm direct binding of TX14A to GPR37L1 using a biotinylated version of the peptide (bTX14A), as described by Meyer et al. (16). Membranes from cells expressing either wild-type or Δ122 GPR37L1-eYFP were incubated with bTX14A before isolation with streptavidin-coated beads. However, no GPR37L1 was evident after pull-down in either these cells (**Figure 6c**) or when we performed the same experiment on endogenously expressed GPR37L1 in mouse cerebellum (**Figure 6d**). Thus, we find no evidence that TX14A binds to GPR37L1.

## Discussion

The key finding of this study is that the orphan GPCR, GPR37L1, is subject to N-terminal proteolysis by matrix metalloproteases both in cultured cells and rodent cerebellum, and that this post-translational modification is linked to loss of Gαs signal transduction. Using various biochemical and pharmacological approaches, we found that GPR37L1 proteolysis likely occurs between the single N-terminal glycosylation site, Asn^105^, and the start of the first transmembrane domain. N-terminal truncation before this putative cleavage site prevented GPR37L1 expression, while truncation of the entire N-terminus (Δ122 GPR37L1) did not alter receptor cell-surface trafficking. Using a yeast chimeric Gpa1/Gα expression system, we also found that full length GPR37L1 displays ligand-independent coupling to Gpa1/Gαs; a response that is lost with the Δ122 GPR37L1 construct. Intrigued by the observation that three different fragments of the GPR37L1 N-terminus have been identified in human CSF (17-19), we posited that GPR37L1 may function in a manner akin to the PARs, which have a tethered agonist that is only revealed upon proteolytic cleavage of the receptor N-terminus (39). However, treatment of GPR37L1 or Δ122 GPR37L1 Gpa1/Gαs yeast transformants with either synthetic peptides corresponding to those identified from human CSF, or the putative tethered peptides that would have remained, does not alter GPR37L1 signaling. Finally, we demonstrated that GPR37L1 proteolytic processing also occurs *in vivo* and that genetic ablation of the receptor reduces cAMP accumulation in *ex vivo* cerebellar cultures, supporting the idea that GPR37L1 can couple to Gαs to stimulate cAMP production.

GPR37L1 is not alone in having its N-terminus processed by a metalloprotease; other examples include the endothelin-B receptor (31), β_1_-adrenoceptor (32), and GPR37 (40)(published during the course of our own study), for which N-terminal cleavage influences cell surface abundance but not activity. By combining multiple processes of elimination, we have narrowed down both the likely mediators of GPR37L1 proteolysis and the N-terminal cleavage site. Few specific ADAM or MMP inhibitors exist, thus we took advantage of the overlapping, but not identical, inhibitory profiles of various small molecule antagonists and TIMPs to identify ADAMs 8, 9, 15, 19, 20, 21, and 30 as possible mediators of GPR37L1 N-terminal truncation; genetic tools will likely be required to directly identify the specific ADAM(s) mediating GPR37L1 N-terminal cleavage. We also sought to identify the metalloprotease cleavage site within the GPR37L1 N-terminus by employing alanine scanning mutagenesis but did not find a mutation that altered proteolysis. This is not unexpected; there is no clear consensus sequence for ADAM or MMP cleavage (36) and amino acid substitutions both within and at variable proximity to the cleaved residues are well tolerated (41). Furthermore, a similar study of N-terminal proteolysis was also unable to prevent receptor cleavage by mutagenesis for the closely related GPR37 (40). Thus, the precise site of GPR37L1 N-terminal cleavage and the identity of the specific protease(s) responsible remain undefined.

Using chimeric yeast Gpa1 fused to the last five amino acids of each human Gα subunit, we provided evidence for GPR37L1 constitutive activity and showed that the receptor couples to Gpa1/Gαs. This may explain the preliminary report of a deleterious effect of adenoviral-mediated GPR37L1 overexpression in cardiomyocytes (12), because it has been reported that excessive β-adrenoceptor stimulated cAMP accumulation decreases cardiomyocyte viability (42), although, unlike the β-adrenoceptor, GPR37L1 protein is undetectable in mouse heart (8), so its heterologous overexpression in cardiomyocytes might also result in non-specific toxicity. The yeast assay relies upon chimeric G proteins and therefore may not report the *in vivo* coupling preference of GPR37L1. Indeed, the constitutive G protein coupling of this receptor in mammalian cells is uncertain: Foster et al. (43) found no evidence for constitutive GPR37L1 activity in any pathway tested (Gαs, Gαi and Gαq), while constitutive Gαi signaling has been reported in human renal proximal tubule cells (44). Moreover, other investigators have reported both constitutive signaling through Gαs (11) and ligand-dependent signaling through Gαi (16). Based upon its amino acid sequence, GPR37L1 is confidently predicted to couple to Gαs (and Gαq) and not to Gαi according to a machine learning-based prediction method, PRECOG (45), that was developed using a comprehensive screen of G protein coupling signatures for 144 Class A GPCRs (46).

Unlike full length GPR37L1, the N-terminal truncation mutant, Δ122 GPR37L1, showed no constitutive signal through Gpa1/Gαs, suggesting that the N-terminus is important for receptor activity and that post-translational modification silences the receptor. For this reason, we tested whether GPR37L1 was activated by a tethered ligand as observed for the PARs, or by the human CSF peptides we had identified from the literature (17-19). That we saw no effect is perhaps unsurprising, because the metalloprotease-mediated processing of GPR37L1 that occurs in cultured cells corresponds to removal of the entire N-terminus, whereas the N-terminal peptides found in human CSF are fragments upstream of this cleavage site. Rather, the human CSF peptides may be indicative of further proteolytic breakdown of the already released GPR37L1 N-terminus, supported by the fact that the prolyl carboxypeptidase cleavage of CSF3 could only occur if a peptide one residue longer than CSF3 existed (19). So, what might be the role of the GPR37L1 N-terminus? Perhaps GPR37L1 constitutive coupling to Gpa1/Gαs results from auto-agonism from the full length N-terminus, as reported for the melanocortin-4 receptor (MC_4_R) (47), which has both an endogenous agonist and inverse agonist (47-49). Or perhaps GPR37L1 does not need a ligand at all, as was recently demonstrated for the closely related orphan GPCR, bombesin 3 receptor, which is also constitutively active (50).

Whether GPR37L1 already has an identified endogenous ligand remains contentious. Using HEK293T cells and primary astrocytes, Meyer et al. (16) reported that neuropeptides TX14A and prosaposin are agonists for both GPR37 and GPR37L1, signaling through Gαi and extracellular signal-regulated kinases 1 and 2 (ERK1/2). However, disentangling the specificity of TX14A/prosaposin for GPR37L1, GPR37 or neither, is problematic as many studies evaluating the potential agonist activity of this ligand have used models in which both receptors are inactivated. For example, Liu et al. (51) showed that simultaneous knockdown of both GPR37 and GPR37L1 in astrocytes abolished TX14A-mediated cAMP inhibition and protection from oxidative toxicity (although intriguingly, they could not replicate TX14A activation of GPR37 or GPR37L1 in HEK293 cells, the same model used in the original discovery of the pairing (16).

Likewise, ERK1/2 phosphorylation in astrocytes was only evident in GPR37 or dual GPR37/GPR37L1 depleted astrocytes (16). While TX14A-mediated inhibition of D-aspartate-evoked current is lost in GPR37L1^-/-^ mice, TX14A-mediated neuroprotection from chemical ischemia was preserved (9). In contrast, our finding that TX14A did not activate or trigger post-translational modification of GPR37L1 is in keeping with several other studies: Southern et al. (52) saw no β-arrestin recruitment to GPR37L1 upon treatment with prosaposin; Giddens et al. (11) saw no change in either cAMP or pERK1/2 levels upon TX14A stimulation despite seeing constitutive activity through these pathways (agreeing with our Gαs observation here); while Foster et al. (43) failed to detect GPR37L1 stimulation by TX14A using β-arrestin recruitment, internalization, and dynamic mass redistribution assays. Finally, we could not reproduce the direct interaction seen between biotinylated TX14A and GPR37L1 reported by Meyer et al. (16). Thus, if TX14A does bind to GPR37L1 it must be with very low affinity and efficacy.

An exciting finding of this study is that N-terminal processing occurs *in vivo* and in cerebellar cultures. In heterologous FlpIN HEK293 and U87 cells, we observed the larger molecular weight GPR37L1 species to be greater or equal in proportion to the smaller, cleaved species. However, our data suggested that, *in vivo*, the proteolytically-processed GPR37L1 predominates. Given our finding that GPR37L1 lacking the N-terminus was inactive in yeast expressing Gpa1/Gαs, the abundance of the proteolyzed form suggests that GPR37L1 may largely exist in an inactivated form due perhaps to very rapid proteolysis *in vivo*. Some signaling-competent receptor must nevertheless be present, because GPR37L1-null mice exhibited reduced cAMP accumulation compared to their wild-type counterparts. Furthermore, changes to extracellular metalloprotease activity would be expected to ‘tune’ basal GPR37L1 signaling by post-translational modification. Future studies will need to examine the role of the individual GPR37L1 protein species in the entire animal to understand the physiological importance of such proteolysis, and to determine whether the cleaved receptor genuinely is signaling incompetent *in vivo*.

In summary, we provide evidence that GPR37L1 is subject to N-terminal proteolysis both *in vitro* and *in vivo*, and that this blunts constitutive Gpa1/Gα_s_ coupling of the receptor in yeast. For many years, orphan GPCRs have been considered an exciting, albeit untapped, resource for new therapeutic targets (53), yet an understanding of the physiology of these receptors or their involvement in disease has been an intractable problem in the absence of a ligand. Here, we show that knowing the ligand is not always necessary for investigating the basic function of a GPCR. Instead, our data and methods revealed that proteolysis may be a mechanism to inactivate a GPCR that may have constitutive activity *in vivo*, because it either does not require a ligand or its ligand is always present and bound to the receptor.

## Methods

### Reagents and constructs

All reagents were from Sigma-Aldrich or Life Technologies (including Invitrogen), unless otherwise stated. A previously reported (16,54) human GPR37L1 was purchased from Multispan; however, this construct lacked the endogenous GPR37L1 signal peptide and instead had an optimized signal peptide and N-terminal Flag tag, so we reintroduced the original sequence with a Hind III restriction site and Kozak sequence by polymerase chain reaction. All constructs in pcDNA5/FRT/TO or pcDNA3 contain human GPR37L1 with eYFP fused to its C-terminus; human GPR37L1-eYFP (wild type and Δ122) was cloned into p426GPD for *S. cerevisiae* experiments. Constructs were verified by DNA sequencing. Epitope tagging of GPR37L1 at the N-terminus and expression in Flp-In HEK293 cells resulted in the appearance of large vacuoles and subsequent cell death. N-terminal CSF and “tethered” peptides were synthesized by GenScript. Chimeric G protein yeast strains were provided by GSK under material transfer agreement.

### Cell culture and transfection

Stable FlpIN T-REx cells were generated and cultured as previously described (25). For all FlpIN assays, cells were cultured in 10% fetal calf serum on poly-d-lysine–coated (0.1 mg/ml) tissue culture dishes for 24 hours before induction with doxycycline. U87 MG cells were cultured in RPMI media containing 10% fetal calf serum, and transiently transfected using Lipofectamine 3000, as per manufacturer protocol.

### Rodent cerebellar tissue harvest and organotypic brain slice cultures

All animal tissue harvests were performed according to the *Australian Code for the Care and Use of Animals for Scientific Purposes, Eighth Edition* (2013), and were approved by either the Garvan Institute of Medical Research/St Vincent’s Hospital Animal Ethics Committee project 13/30 (mouse) or the University of New South Wales Animal Care and Ethics Committee project 15/12B (rat). GPR37L1-null mice were originally described in Min et al. (12) and reanimated from sperm by Charles River Laboratories (Japan). C57BL/6J wild-type or GPR37L1-null male mice between 4 and 6 months of age were euthanized by cervical dislocation and cerebella were isolated and snap-frozen in liquid nitrogen for −80°C storage until required. Organotypic cerebellar slice cultures were prepared essentially as described for the hippocampus (55,56). Briefly, 15-day-old mixed-sex, C57BL/6J wild-type or GPR37L1-null mice or 8- to 9-day-old mixed-sex Sprague Dawley rats were anaesthetized and then euthanized by decapitation, and brains were removed and placed in ice-cold Hanks’ balanced salt solution (HBSS) supplemented with 30 mM d-glucose and penicillin/streptomycin (100 U/ml) and saturated with carbogen (95% oxygen, 5% carbon dioxide). The cerebella were removed and sliced into 400 μm sagittal sections using a McIlwain tissue chopper and then plated evenly across two Millicell cell culture inserts (Merck) in six-well plates (two inserts per cerebellum, six to eight slices per insert). All rodent cultures were maintained in culture media containing 50% minimum essential medium, 25% HBSS, 25% heat-inactivated horse serum, 36 mM d-glucose, 25 mM HEPES, 4 mM NaHCO_3_, 2 mM GlutaMAX, 1% Fungizone, and penicillin/streptomycin (100 U/ml). Rat cerebellum cultures were maintained at 37°C in an atmosphere of 5% CO_2_, and culture media were changed every 3 days. After 10 days in culture, a single well of each matched pair was treated by an addition of 20 μM BB94 to culture media (the other well in each pair was left untreated). Forty-eight hours after treatment, slices were collected and homogenized in ice-cold RIPA (radioimmunoprecipitation assay) buffer, as described below. Mouse cerebellar cultures were grown in identical conditions but were harvested after 1 day in culture, with or without 20 or 40 μM BB94. Data were obtained from at least three independent litters of animals.

### In-gel fluorescence and immunoblot studies

After GPR37L1 induction in FlpIN cells by doxycycline for 24 hours (or after 24 h of transient transfection in the case of U87 MG cells), cells were placed on ice, washed with ice-cold phosphate-buffered saline, and harvested with ice-cold RIPA buffer [50 mM Hepes, 150 mM NaCl, 1% Triton X-100, 0.5% deoxycholate, 0.1% SDS, 10 mM NaF, 5 mM EDTA, 10 mM NaH_2_PO_4_, 5% ethylene glycol, and cOmplete EDTA-free Protease Inhibitor Cocktail Tablet (pH 7.4) (Roche Diagnostics)]. Cell lysates were cleared of insoluble debris by centrifugation, and the supernatant was mixed with Laemmli buffer [final concentration was as follows: 63 mM Tris, 50 mM dithiothreitol, 80 mM SDS, 10% glycerol, 0.004% bromophenol blue (pH 6.8)], followed by 15 min incubation in a 37°C water bath. For deglycosylation experiments, cleared lysates were treated with either 2 U of peptide *N*-glycosidase F or 100 mU of endoglycosidase H, or both, for 2 hours in a 37°C water bath before addition of Laemmli buffer. Cell surface biotinylation was performed as previously described (57). Equal amounts of protein (as determined by Merck Direct Detect) were then separated by 4 to 12% bis-tris SDS-PAGE in NuPAGE MOPS SDS Running Buffer (Life Technologies) either at room temperature (for Western blot) or at 4°C in ice-cold buffer [for in-gel fluorescence, imaged with a FLA-5100 (Fujifilm)]. For immunoblotting, proteins were transferred to a 0.45 μm pore size polyvinylidene fluoride transfer membrane (Merck Millipore), and nonspecific binding was blocked in Tris-buffered saline/0.1% Tween 20 with 5% skim milk powder. GPR37L1-eYFP was detected with an antibody against GFP (1:10,000 dilution; Abcam) and IgG anti-rabbit horseradish peroxidase–conjugated secondary antibody (1:10,000; GE Healthcare UK). For whole cerebellum and cerebellar slices, snap-frozen tissue was prepared using a POLYTRON homogenizer in the presence of RIPA buffer, and insoluble debris was removed as described above. Crude membranes were isolated as previously described (26). Endogenous GPR37L1 was detected by goat anti-GPR37L1 (C-12) antibody (1:1000; sc-164532, Santa Cruz Biotechnology). Chemiluminescence was detected with Western Lightning ECL reagent (PerkinElmer).

### Inhibitor studies and verification of recombinant TIMP activity

Metalloprotease inhibitors, including BB94 (batimastat), TAPI-1 and TAPI-2 (Santa Cruz Biotechnology), and TIMP1, TIMP2, TIMP3, and TIMP4 (R&D Systems), were added at the time of GPR37L1 induction (or 5 hours after transfection in the case of U87 MG cells) or 24 hours after induction for TX14A experiments, and cells were then incubated for 24 hours before lysis. For MG132 proteasome inhibition, cells were induced with doxycycline overnight and then treated with MG132 for 6 hours before harvest. Verification of TIMP activity *in vitro* using activated MMP-2 and the fluorogenic peptide substrate MCA-Pro-Leu-Gly-Leu-DPA-Ala-Arg-NH_2_ was performed as per the protocol provided by R&D Systems.

### Assessment of G protein coupling using S. cerevisiae G protein chimeras

p426GPD (high-copy vector), p426GPD–GPR37L1-eYFP, or p426GPD–Δ122 GPR37L1-eYFP was transformed into 11 individual FUS1-regulated β-galactosidase yeast reporter strains that had been modified to express either the endogenous Gα protein Gpa1 or chimeras comprising Gpa1/Gα fusions containing the last five amino acids of each human G protein (22). Individual *S. cerevisiae* transformants were then screened for constitutive coupling through yeast chimeras using the same protocol as previously described (21). Fluorescence was detected using a BMG PHERAstar FS (excitation/emission, 485/520, in AFU) (BMG).

### cAMP ELISA

To assess the cAMP levels in mouse cerebellar slices, tissue was harvested as described above, except that slices were incubated for exactly 1 hour in the presence or absence of the phosphodiesterase inhibitor IBMX (1 mM). To ensure that there was sufficient tissue for the assay, slices from two to four separate mixed-sex pups were pooled and considered to represent *n* = 1. cAMP was measured using the DetectX Direct cAMP ELISA (enzyme-linked immunosorbent assay) kit (Arbor Assays), as per manufacturer’s instructions.

### Data analysis

All data analysis was performed with GraphPad Prism 6 (GraphPad Software Inc.). Graphs show means ± SEM. Where variances were unequal according to a Brown-Forsythe test, and therefore violating the assumptions of ANOVA, data were transformed to logarithmic values before one-way ANOVA analysis. In-gel fluorescence densitometry was performed using ImageJ software (National Institutes of Health) within the linear range, and immunoblots were processed without altering relative intensities of the bands.

### Data and materials availability

Yeast chimeras are available from GlaxoSmithKline under an MTA. All other materials are available from the authors upon request.

## Acknowledgements

We thank Dr Angela Finch (UNSW Sydney) for critical reading of the manuscript and helpful suggestions, and Dr Irina Kufareva (UC San Diego) for pointing us towards the glioblastoma model. We also thank Drs Bryan Day and Seçkin Akgül (QIMR Berghofer Medical Research Institute) for providing relevant glioblastoma cell lines and helpful advice. Yeast chimeras were provided by S. Dowell from GlaxoSmithKline under a material transfer agreement (MTA).

This work was funded in part by National Health and Medical Research Council of Australia (NHMRC) Program Grants 573732 and 1074386 (R.M.G.), a NHMRC/National Heart Foundation of Australia (NHF) C.J. Martin Fellowship and NHF Future Leader Fellowship to N.J.S., Australian Postgraduate Awards (J.L.J.C. and T.N.), and a Simon and Michal Wilkenfeld Scholarship (J.L.J.C.).

## Author Contributions

N.J.S. designed the study, supervised the project, performed the experiments, and wrote the manuscript. J.L.J.C. and T.N. performed the experiments and wrote the manuscript. R.E.S. and A.J.C. performed experiments with U87 cells. N.M.J. designed and performed organotypic cerebellar slice assays with J.L.J.C.. R.M.G. co-supervised the project and provided critical input. All authors have contributed to the editing of the manuscript.

## Footnote

The studies described herein have been previously published in a manuscript that was retracted at the authors’ request after an error was discovered in one of the constructs (Coleman et al. (58), *Metalloprotease cleavage of the N terminus of the orphan G protein-coupled receptor GPR37L1 reduces its constitutive activity*. Sci Signal. 9, ra36). None of the affected data has been included in the current study and the remaining work is presented here with permission from AAAS.

## Supplementary Figure

**Supp 1:**
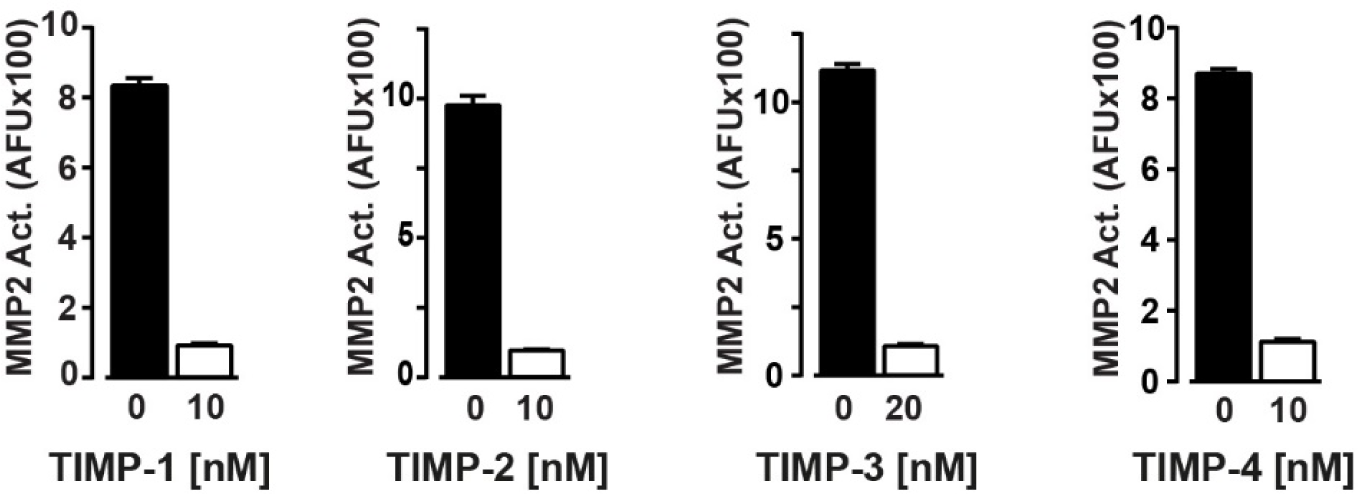
Human recombinant Tissue Inhibitors of Metalloproteases were biologically active. Human recombinant TIMP-1, TIMP-2, TIMP-3, or TIMP-4 biological activity was verified *in vitro* using activated MMP2 and the fluorogenic substrate, MCA-Pro-Leu-Gly-Leu-DPA-Ala-Arg-NH_2_, as per the protocol provided by R&D Systems. AFU: arbitrary fluorescence units.

